# Neutrophils infiltrate sensory ganglia and mediate chronic widespread pain in fibromyalgia

**DOI:** 10.1101/2022.06.29.498149

**Authors:** Sara Caxaria, Sabah Bharde, Alice M. Fuller, Romy Evans, Bethan Thomas, Petek Celik, Francesco Dell’Accio, Simon Yona, Derek Gilroy, Mathieu-Benoit Voisin, John N. Wood, Shafaq Sikandar

**Author notes:** Corresponding authors: Shafaq Sikandar and John N Wood.

## Abstract

Fibromyalgia is a debilitating widespread chronic pain syndrome that occurs in 2-4% of the population. The prevailing view that fibromyalgia results from central nervous system dysfunction has recently been challenged with data showing changes in peripheral nervous system activity. Using a mouse model of chronic widespread pain through hyperalgesic priming of muscle, we show that neutrophils invade sensory ganglia and confer mechanical hypersensitivity on recipient mice, whilst adoptive transfer of immunoglobulin, serum, lymphocytes or monocytes have no effect on pain behaviour. Neutrophil depletion abolishes the establishment of chronic widespread pain in mice. Neutrophils from patients with fibromyalgia also confer pain on mice. A link between neutrophil derived mediators and peripheral nerve sensitisation is already established. These observations suggest new approaches for targeting fibromyalgia pain through an understanding of the mechanisms that cause altered neutrophil activity and interactions with sensory neurons.

**Significance statement:** We used a back-translational model in mice to demonstrate the pro-nociceptive role of neutrophils in fibromyalgia. Adoptive transfer of neutrophils from mice with chronic widespread pain or from patients with fibromyalgia can confer mechanical pain to recipient naïve mice, sensitise evoked action potential firing of spinal cord neurons and produce phenotypic changes in cell surface expression of neutrophil proteins that cause infiltration of neutrophils into dorsal root ganglia. These data provide the framework for an immunological basis of chronic widespread pain in fibromyalgia mediated by polymorphonuclear granulocytes.

## Introduction

Although the aetiology of chronic widespread pain in fibromyalgia syndrome is unknown, the concept of altered central processing of nociceptive information has dominated the literature and clinical treatment guidelines (1). Non-specific analgesics that modulate monoaminergic pathways in the central nervous system are conventionally used for treating the maladaptive plasticity that accompanies persistent, widespread hypersensitivity in patients diagnosed with fibromyalgia syndrome. However, several studies point towards a peripheral drive of chronic widespread pain in fibromyalgia syndrome, suggesting that a targeted approach towards a peripheral aetiology is possible. For example, in patients with fibromyalgia syndrome, peripherally administration of a localized lidocaine block of tender points reduces pain scores (2). Interestingly neutrophil function as well as nociceptor activity involves lidocaine-sensitive sodium channels (3). Moreover, abnormal activity-dependent slowing of unmyelinated nociceptors is observed in microneurography recordings (4); and pain-related evoked potentials reflect abnormalities in small fibre function (5).

Aberrant activity of immune cells and associated cytokine signaling has also been linked to fibromyalgia syndrome. Neutrophils are polymorphonuclear granulocytes that normally exist in circulation to function as primary mediators of rapid innate host defense, but are surprisingly found in higher number in the circulation of fibromyalgia patients, with a phenotype characterized by enhanced chemotactic and microbicidal properties (6-8). Systemic expression of proinflammatory cytokines, such as IL-6, IL-8 and TNF-α, is increased in patients with fibromyalgia, and these same cytokines are otherwise released by neutrophils and are known mediators of nociceptor sensitisation (9-11).

Growing evidence demonstrates an intricate bidirectional interaction between immune cells and sensory neurons. Polarisation of resident macrophages in dorsal root ganglia and mitochondrial transfer from infiltrating macrophages to somata can confer sensitisation of peripheral sensory neurons (12, 13). Similarly, CD3+ T-cells have also been described in the resolution pathway of inflammatory pain (14) and pharmacological blockade of T cell-derived leukocyte elastase significantly reduces the magnitude of behavioural hypersensitivity in a rodent model of neuropathic pain (15). Conversely, Na_v_1.8+ sensory neurons play a key role in psoriasis and CD8+ T-cell responses to viral infection (16). Neutrophils are not normally resident cells within intact nervous tissue, but infiltration into peripheral nerves has been reported in rodent models of nerve injury (17, 18), diabetic neuropathy (19) and degenerative disease (20).

Here we investigate a causal link between neutrophils and nociceptive activity underlying chronic widespread pain in fibromyalgia using a mouse model of hyperalgesic priming (21-23) and adoptive transfer of cells from mice and patients with chronic pain to recipient naïve mice. Integrating a combination of pain behaviour, electrophysiological measures of neuronal excitability and imaging of peripheral sensory ganglia, we demonstrate pro-nociceptive activity of neutrophils.

## Results

### Persistent hypersensitivity in a mouse model of chronic widespread pain

A model of chronic widespread pain was induced in male mice using a hyperalgesic priming paradigm to trigger latent and long-lasting pain behaviour with consecutive intramuscular injections of carrageenan, an acute inflammatory stimulus (Figure 1A) (23). We compared pain behaviour between primed mice and control mice, where the latter group only received one intramuscular injection to induce acute inflammatory pain. Both Primed and Control mice developed mechanical hypersensitivity to von Frey stimulation of the hind paw after the first carrageenan administration that resolved by 4d (Figure 1Bi; 0.91<u>±</u>0.1g to 0.38<u>±</u>0.1g at 1d in control mice; 1.2<u>±</u>0.2g to 0.37<u>±</u>0.1g at 1d in primed mice). Primed mice developed persistent ipsilateral mechanical hypersensitivity to von Frey stimulation following the second intramuscular injection of carrageenan, with later development of contralateral hypersensitivity by ca. 2 weeks after induction of the model. Although Control mice developed transient ipsilateral hypersensitivity after the first injection on D0 (Fig. 1Bi), they did not develop any prolonged ipsilateral or contralateral hypersensitivity for the duration of the study. Mechanical withdrawal thresholds throughout the remaining duration of the experiment were lower in both ipsilateral and contralateral paws in Primed mice compared to Control mice (Figure 1Bii). Control mice and Primed mice developed weightbearing asymmetry after the first injection on D0 compared to respective baselines, but Primed mice had sustained weightbearing asymmetry compared to control mice until D12, when contralateral mechanical hypersensitivity had developed in Primed mice, resulting in equal weight bearing of both hindlimbs (Figure 1C). In contrast to the development of mechanical hypersensitivity, heat pain thresholds of Primed mice were comparable to control mice in the hotplate and Hargreaves’ assays for the duration of the hyperalgesic priming paradigm (Figure 1D and Figure 1E). Moreover, priming did not affect motor function measured in the rotarod (Figure S1).

**Figure 1:**
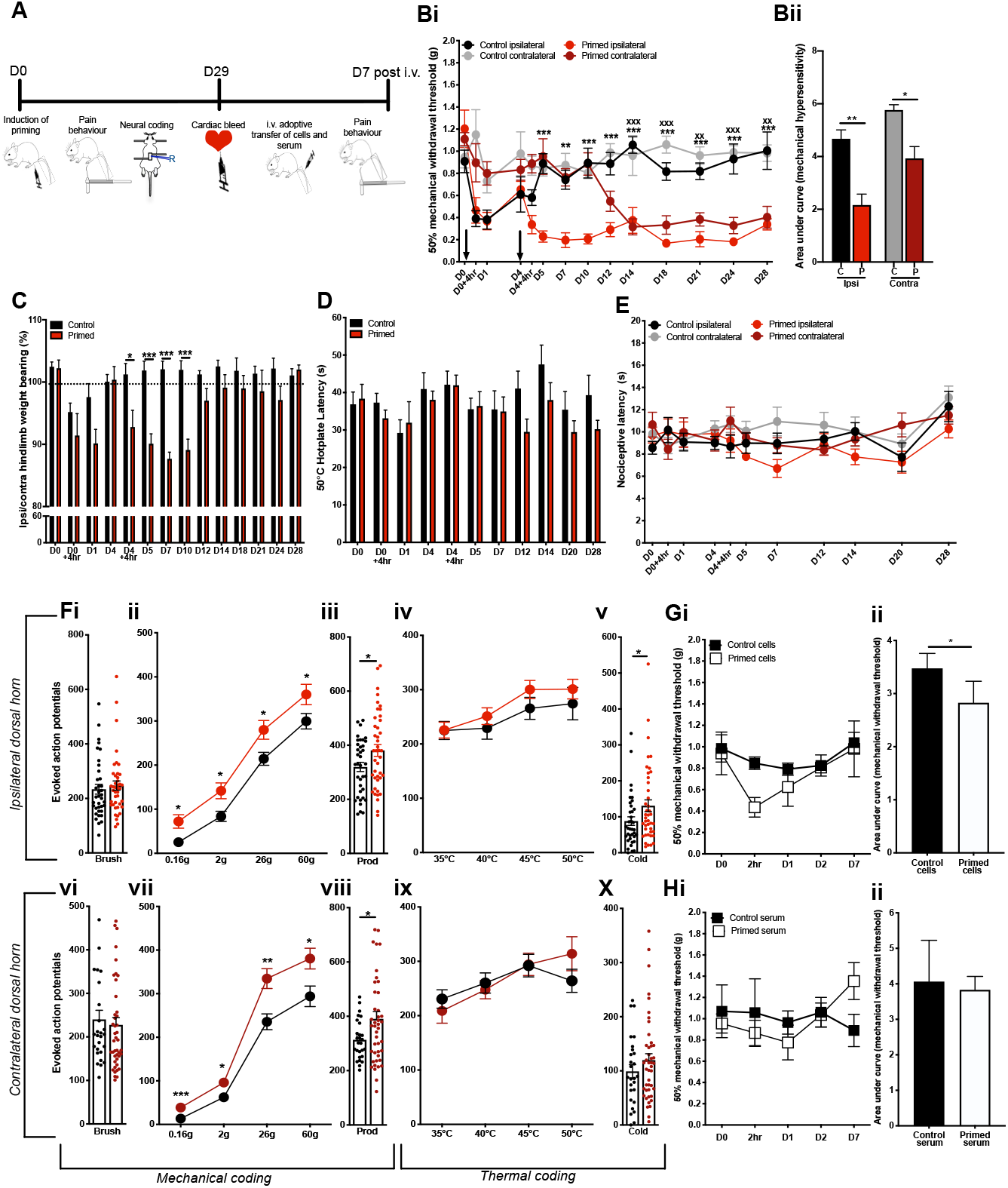
A hyperalgesic priming model for widespread pain in mice. **(A)** Schematic for study design: a hyperalgesic priming model of chronic widespread pain was induced in mice using intramuscular injections of carrageenan on D0 and D4 in Primed mice (*n*=7). Control mice (*n*=7) received a single intramuscular injection on D0. After D29 cells and serum were obtained following cardiac bleed from Primed and Control mice and injected into recipient naïve mice whose pain behaviour was monitored up to 7 days. **(Bi)** Mechanical hypersensitivity to von Frey stimulation of ipsilateral and contralateral limbs in Primed mice (red) and Control mice (**P<0.01, ***P<0.001 2 way RM ANOVA Primed ipsilateral vs Control ipsilateral; ^^P<0.01, ^^^P<0.001 2 way RM ANOVA Primed contralateral vs Control contralateral). **(Bii)** Area under curve of mechanical pain thresholds illustrated in (Bi) of ipsilateral and contralateral hindlimbs in Primed and Control mice (*P<0.05, **P<0.01 Unpaired t-test). **(C)** Weightbearing asymmetry of hind limbs (*P<0.05, ***P<0.001 2 way RM ANOVA Primed vs Control). **(D)** Latency to nociceptive behaviour in the hotplate assay (2 way RM ANOVA Primed vs Control). **(E)** Latency to nociceptive withdrawal in Hargreaves’ assay (2 way RM ANOVA Primed ipsilateral vs Control ipsilateral, Primed contralateral vs Control contralateral). **(F)** Evoked action potential firing of neurons in the (i-v) ipsilateral dorsal horn (control cells n=37, n=42 primed cells) and in the (vi-x) contralateral deep dorsal horn (control cells n=27, n=44 primed cells) of Primed mice and Control mice to (i, vi) innocuous brush stimulation, (ii, vii) von Frey stimuli, (iii, viii) noxious prod stimulation, (iv, ix) heat stimuli and (v, x) noxious cold stimulation with ethyl chloride (*P<0.05, **P<0.01 Unpaired t-test). **(G)** Mechanical hypersensitivity following adoptive transfer of blood cells from primed mice (n=4) and control mice (n=4) (*P<0.05 Unpaired t-test). **(H)** Mechanical hypersensitivity of naïve mice following adoptive transfer of blood serum from Primed mice (n=4) and Control mice (n=4) (Unpaired t-test).

We performed multi-unit extracellular recordings in the dorsal horn of live mice to determine how the induction of widespread pain in the hyperalgesic priming model affects the bilateral excitability of spinal cord neurons, which are otherwise responsible for mediating pain-related reflexes in the pain behaviour assays we used. We quantified the number of action potentials fired by polymodal wide dynamic range neurons in the ipsilateral and contralateral dorsal horn in response to 10s of peripheral stimulation of various modalities (Figure 1F). Ipsilateral and contralateral dorsal horn neurons in Control and Primed mice displayed graded intensity coding to innocuous and noxious mechanical punctate stimulation with von Frey hairs, although Primed cells displayed a sensitised response and consistently fired more action potentials to all von Frey intensities (Figure 1Fii, 1vii). Dorsal horn cells in Primed mice were also sensitised to noxious prod stimulation of the hindpaw compared to cells in Control mice (Figure 1Fvii-viii). Dorsal horn cells in Primed mice did not show any significant difference in evoked activity to brush or heat stimuli (Figure 1Fi,iii, vi, ix). These findings support our observation that Primed mice did not develop behavioural hypersensitivity to noxious heat stimulation, and further suggest that induction of the hyperalgesic priming model does not confer sensitisation to innocuous dynamic mechanical stimulation. However, we did observe sensitisation of Primed cells triggered by application of a noxious cold stimulus, which was only statistically significant in ipsilateral cells (average 88<u>±</u>11 spikes in primed cells compared to 131<u>±</u>16 spikes in control cells ipsilateral side; average 99<u>±</u>12 spikes in primed cells compared to 120<u>±</u>11 spikes in control cells contralateral side). Together these data demonstrate that induction of the hyperalgesic priming model leads to the development of persistent pain behaviour encoded by prolonged ipsilateral hypersensitivity at the spinal cord level and a spatiotemporal spread of mechanical hypersensitivity to the contralateral limb.

### Circulating neutrophils are essential for the development of hypersensitivity in a mouse model of chronic widespread pain

We further investigated whether a systemic circulating factor could underlie the observed contralateral hypersensitivity in primed mice. We first transferred isolated blood cells and serum from Primed and Control mice to recipient naïve mice and observed resulting changes in pain behaviour. Transfer of blood cells, but not serum, from Primed mice induced transient mechanical hypersensitivity lasting up to 1d (Figure 1G, Figure 1H). These observations suggest that circulating cells constitute the algogenic aetiology of chronic widespread pain in this murine model.

In order to determine the identity of cells driving pain behaviour following adoptive transfer of blood cells from primed mice, we flow sorted blood cells derived from primed mice displaying persistent widespread hypersensitivity and from control mice with normal pain thresholds (Figure 2A-2C). Four different immune cell types – T-cells, B-cells, neutrophils and monocytes – from Primed mice and Control mice were administered i.v. to naïve recipient mice, and resulting pain behaviours were measured (Figure 2D-E). Mice that were administered neutrophils derived from Primed mice displayed robust, transient mechanical hypersensitivity in the von Frey assay compared to mice that were administered neutrophils derived from control mice (Figure 2Diii). In order to validate the algogenic role of neutrophils in mediating chronic widespread pain, we pharmacologically depleted circulating neutrophils with systemic administration of 1A8 clone on D3, just prior to the second intramuscular injection in primed mice (Figure 3). Administration of the anti-neutrophil antibody did not affect resolution of acute pain behaviour following an intramuscular injection of carrageenan (Figure 3A). However, primed mice administered with this antibody failed to develop prolonged ipsilateral hypersensitivity and any contralateral hypersensitivity compared to control primed mice administered with PBS (Figure 3B). These observations suggest that neutrophil depletion prevents development of persistent and widespread pain, but does not affect gross measures of inflammation and its resolution (Figure 3C-3D). Neutrophil depletion lasted at least 24hr as confirmed by FACS of circulating CD45+ cells, double positive for Ly6G and Ly6C as markers of neutrophils (Figure 3E).

**Figure 2:**
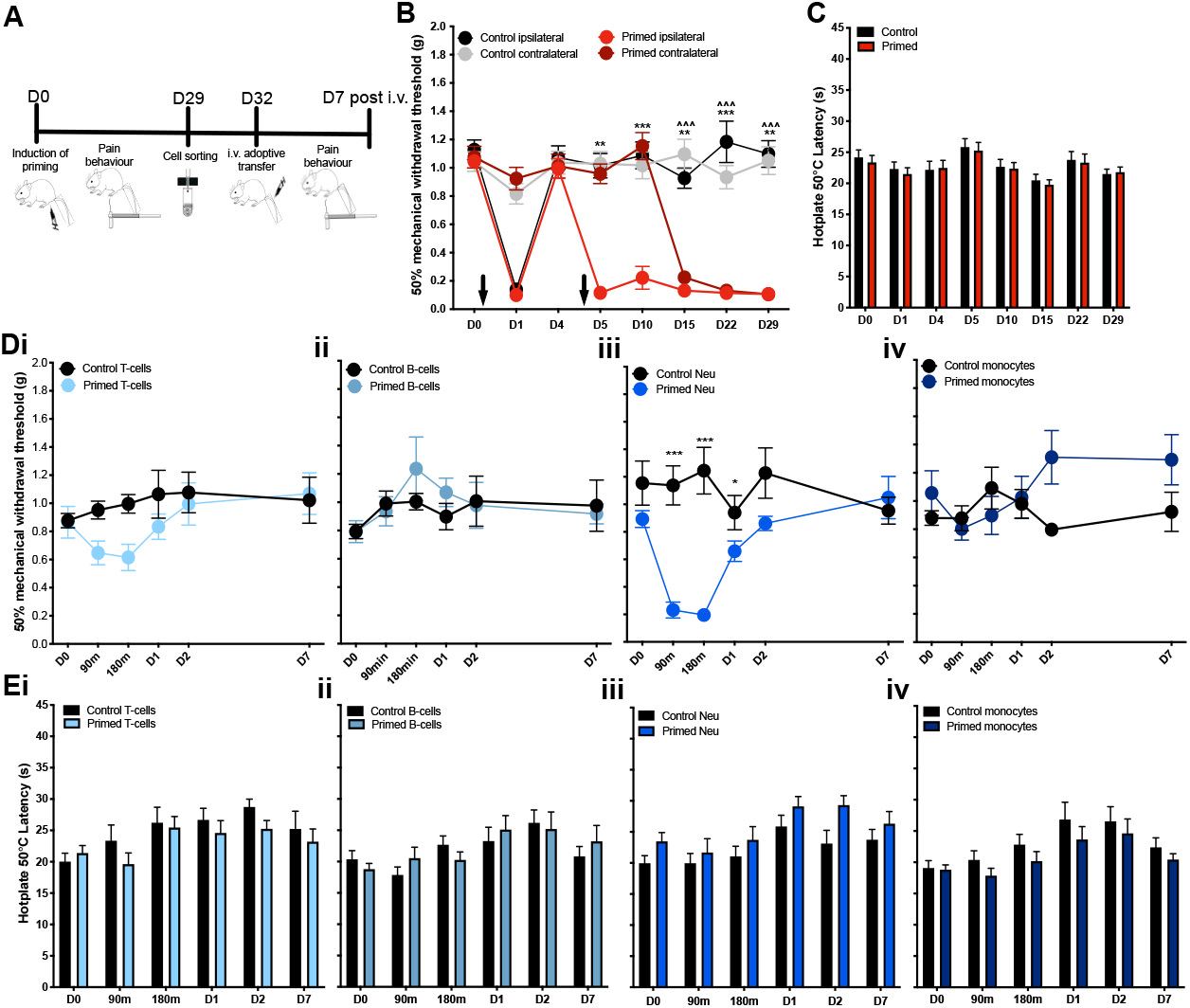
Adoptive transfer of neutrophils from primed mice confers widespread pain in naïve mice. **(A)** Schematic for study design: a hyperalgesic priming model of widespread pain was induced mice (Primed mice n=24; Control mice n=24). On D29 blood cells were obtained through cardiac bleed from Primed and Control mice, flow sorted into B-cells, T-cells, neutrophils and monocytes and injected into naïve mice whose pain behaviour was monitored up to 7 days. **(B)** Mechanical hypersensitivity to von Frey stimulation of ipsilateral and contralateral limbs of Primed and Control mice (**P<0.01, ***P<0.001 2 way RM ANOVA Primed ipsilateral vs Control ipsilateral; ^^^P<0.001 2 way RM ANOVA Primed contralateral vs Control contralateral). **(C)** Latency to nociceptive behaviour in the hotplate assay (2 way RM ANOVA Primed vs Control). **(D)** Mechanical hypersensitivity to von Frey stimulation and **(E)** thermal hypersensitivity in the hotplate following adoptive transfer of (i) T-cells, (ii) B-cells, (iii) neutrophils and (iv) monocytes from primed mice (n=8) and control mice (n=8) (*P<0.05, ***P<0.001 2 way RM ANOVA Primed vs Control cells).

**Figure 3:**
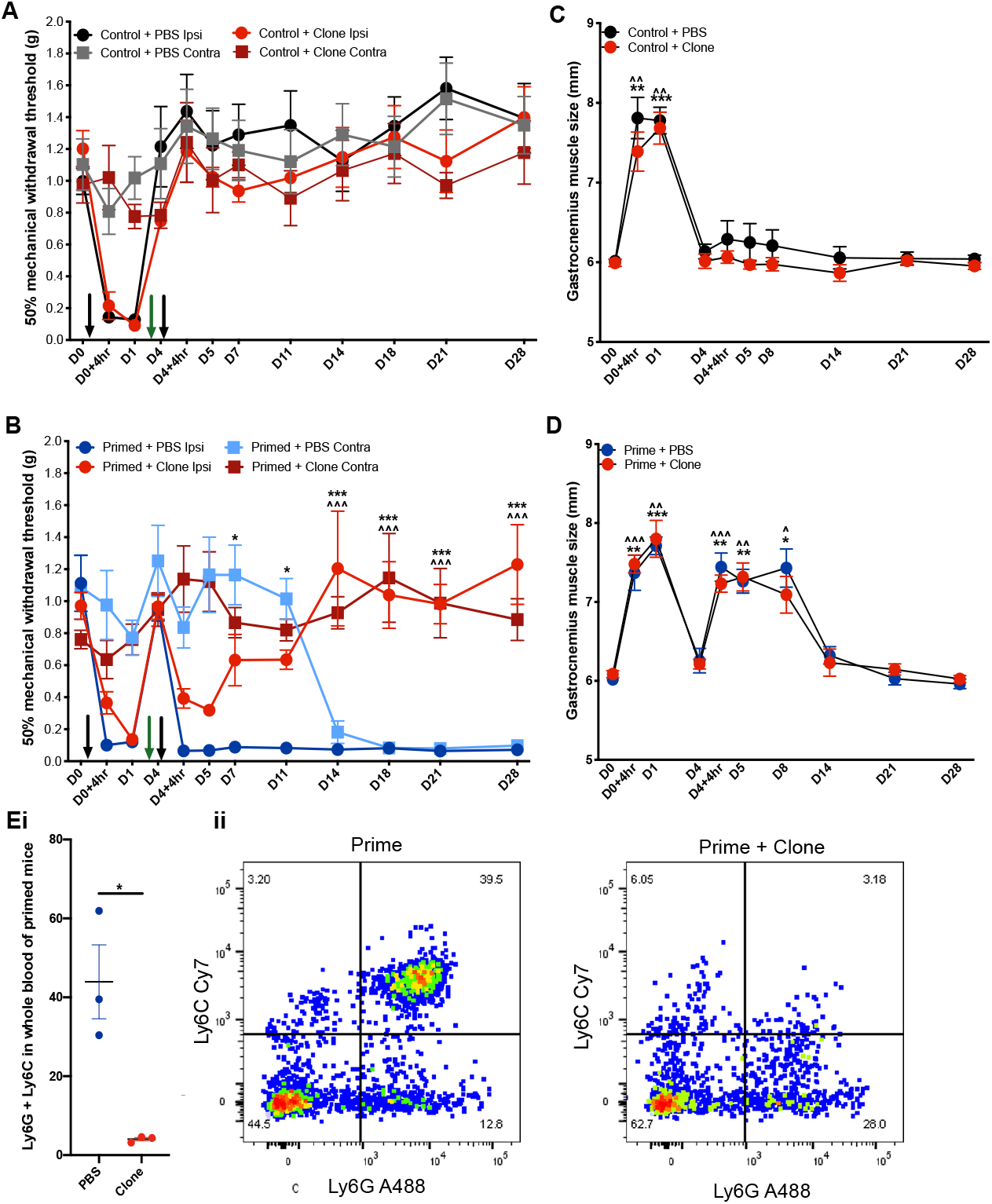
Neutrophil depletion prevents development of widespread pain. A hyperalgesic priming model of widespread pain was induced mice with intramuscular injections of carrageenan on D0 and D4 (black arrows) and neutrophil sequestering antibody (50µg i.v. 1A8 Clone) administered on D3 (green arrow) (Primed + Clone *n*=8; Primed + PBS *n*=8; Control + Clone *n*=8; Control + PBS *n*=8). **(A)** Mechanical hypersensitivity to von Frey stimuli in Control groups receiving one intramuscular injection of carrageenan on D0 and Clone on D3. **(B)** Mechanical hypersensitivity to von Frey stimuli in Primed groups receiving two intramuscular injections of carrageenan on D0 and D4 and Clone on D3 (*P<0.05, ***P<0.001 2-way RM ANOVA Primed + PBS ipsilateral vs Primed + Clone ipsilateral; ^^^P<0.001 2 way RM ANOVA rimed + PBS contralateral vs Primed + Clone contralateral). **(C-D)** Hindleg inflammation measured through gastrocnemius muscle size. **(Ei)** FACS confirmation of neutrophil depletion using 1A8 clone and **(ii)** forward and side scatter for Ly6G and Ly6C markers of neutrophils.

### Neutrophils derived from mice with chronic widespread pain infiltrate sensory ganglia

To test the hypothesis that circulating neutrophils in primed mice have the capacity to interact and cause sensitisation of peripheral sensory neurones, we measured whether neutrophils were directly interacting with somata within dorsal root ganglia. We observed significant *de novo* infiltration neutrophils into L4 DRG of Primed mice compared to Control and naïve animals (Figure 4A). Similarly, we used FACS to confirm greater numbers Ly6G/Ly6C double positive CD45+ cells in L3-L5 DRGs of Primed mice compared to Controls (Figure 4B). We then characterized neutrophils from blood and from DRG in Control and Primed mice and compared the prevalence of subpopulations or ‘clusters’ Ly6G/Ly6C double positive cells using an unsupervised t-distributed stochastic neighbour embedding (tSNE) analysis (Figure S2 and Figure 4C-F). Distinct neutrophil subpopulations from blood and DRG tSNE were defined and clustered by cell surface markers CXCR2, CD62L, CD11b, Ly6G and Ly6C (Figure S2i-ii). We applied tSNE analyses to neutrophil populations derived from Control and Primed mice to identify 11 distinct subpopulations (clusters) in the blood (Figure 4C) and DRG (Figure 4E) based on density of expression. The most upregulated and downregulated clusters of neutrophils derived from blood (Figure 4D) and from DRG (Figure 4F) illustrate differences in the expression of CXCR2 and CD62L, which was further confirmed by independent FACS analysis of Ly6G/Ly6C double positive cells in blood (Figure 4G) and DRG (Figure 4H) of Control and Primed mice. Furthermore, tSNE was performed for all neutrophils (blood and DRG pooled together) to illustrate an enriched neutrophil subpopulation in DRG that is characterized by decreased levels of CXCR2, CD62L, CD11b, Ly6G and Ly6C (Figure S2iii).

**Figure 4:**
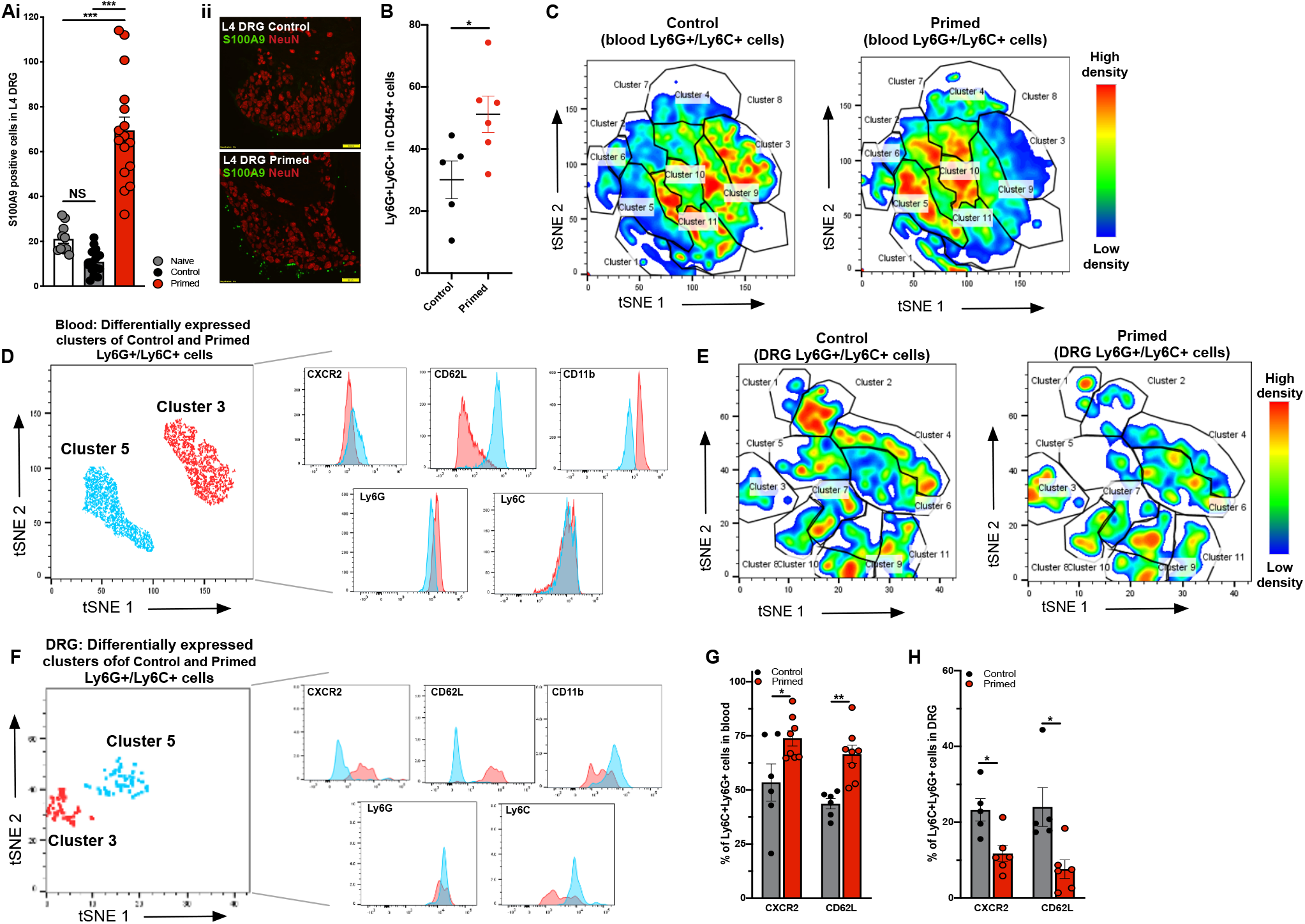
Chronic widespread pain in mice increases neutrophil trafficking into sensory ganglia. **(A)** Immunohistochemical staining for S100A9 in L4 DRG sections from Control and Primed mice **(i)** demonstrating increased numbers of neutrophils in DRG of Primed mice (sections from *n*=3-4 mice per group; ***P<0.001 One-way ANOVA). Yellow line in representative L4 DRG images images **(ii)** indicates 100µm. **(B)** FACS sorted number of neutrophils (Ly6G/Ly6C double positive cells) derived from DRG in Control (*n*=5) vs Primed (*n*=6) mice (*P<0.05 Unpaired T-test). **(C)** tSNE analysis of Ly6C/Ly6G double positive neutrophils obtained from blood of Control (*n*=6) mice and Primed (*n*=8) mice. Neutrophil subpopulations labelled for CXCR2, CD62L, CD11b, Ly6G and Ly6c are illustrated with colours in the expression level heatmap representing median intensity values for relative clustering based on cell density plots. **(D)** Differential expression of CXCR2, CD62L, CD11b, Ly6G and Ly6C based on Cluster 3 and Cluster 5, with the most decreased (−55%) and increased (+101%) numbers of neutrophil subpopulations in Primed cells, respectively. **(E)** tSNE analysis of Ly6C/Ly6G double positive neutrophils obtained from DRG of Control mice (*n*=5) and Primed (*n*=6) mice, with CXCR2, CD62L, CD11b, Ly6G and Ly6c markers. **(F)** Differential expression of CXCR2, CD62L, CD11b, Ly6G and Ly6C based on Clusters 3 and 5, with the most decreased (−60%) and increased (+175%) numbers of neutrophil subpopulations in Primed cells, respectively **(G-H)** FACS quantification of CXCR2 positive and CD62L positive neutrophils (double positive for Ly6G and Ly6C) in **(G)** blood vs **(H)** DRG (*P<0.05, **P<0.01 Unpaired T-test).

### Neutrophils derived from patients with fibromyalgia syndrome induce mechanical hypersensitivity in mice

In order to further evaluate the pro-nociceptive activity of neutrophils, we used a back-translational model of adoptive transfer using neutrophils derived from patients diagnosed with fibromyalgia syndrome administered to naïve mice (Figure 5A). All patients had a minimum of VAS 50 for scoring their pain on the day they were consented to provide blood (mean VAS 68.8<u>±</u>4.2), as well as their overall pain during the week (mean VAS 78.8<u>±</u>3.6). healthy subjects were largely pain free (mean VAS for the day 0.8<u>±</u>0.5 and mean VAS for the week 1.4<u>±</u>0.9) (Figure 5B). Neutrophils derived from patients with widespread pain conferred robust mechanical hypersensitivity in recipient naïve mice compared to neutrophils from healthy control subjects (Figure 5C), but did not affect thermal pain thresholds (Figure 5D). In order to determine effects on neuronal excitability, we measured the effects of adoptive neutrophil transfer on neural coding of wide dynamic range deep dorsal horn neurons (Figure 5E). Spinal cord cells of naïve mice demonstrated marked sensitization to noxious mechanical stimulation of receptive fields in the hindpaw with von Frey hairs, prod and pinch, within 1hr of administration of neutrophils (Figure 5Eiii, 5Eiv), but no marked change in coding of innocuous dynamic stimulation with brush (Figure 5Ei). In addition, we observed sensitisation of dorsal horn neurons to noxious cold stimulation, and surprisingly, also noxious heat stimulation with 50°C despite the lack of effect on 50°C hotplate-evoked behaviours (Figure 5D and 5Eii). We used *ex vivo* imaging of L4 ganglia derived from naïve mice 24hr following i.v. administration of neutrophils from either a patient with fibromyalgia (P) or pain-free control subject (C) and observed greater numbers of neutrophils infiltrating ganglia following adoptive transfer of patient neutrophils (Figure 5F). The neutrophils infiltrating sensory ganglia were both endogenous to the mouse and exogenous human neutrophils, suggesting tissue-restricted innate mechanisms for the recruitment of endogenous and exogenous neutrophils into nervous tissue in chronic pain states (Figure S3).

**Figure 5:**
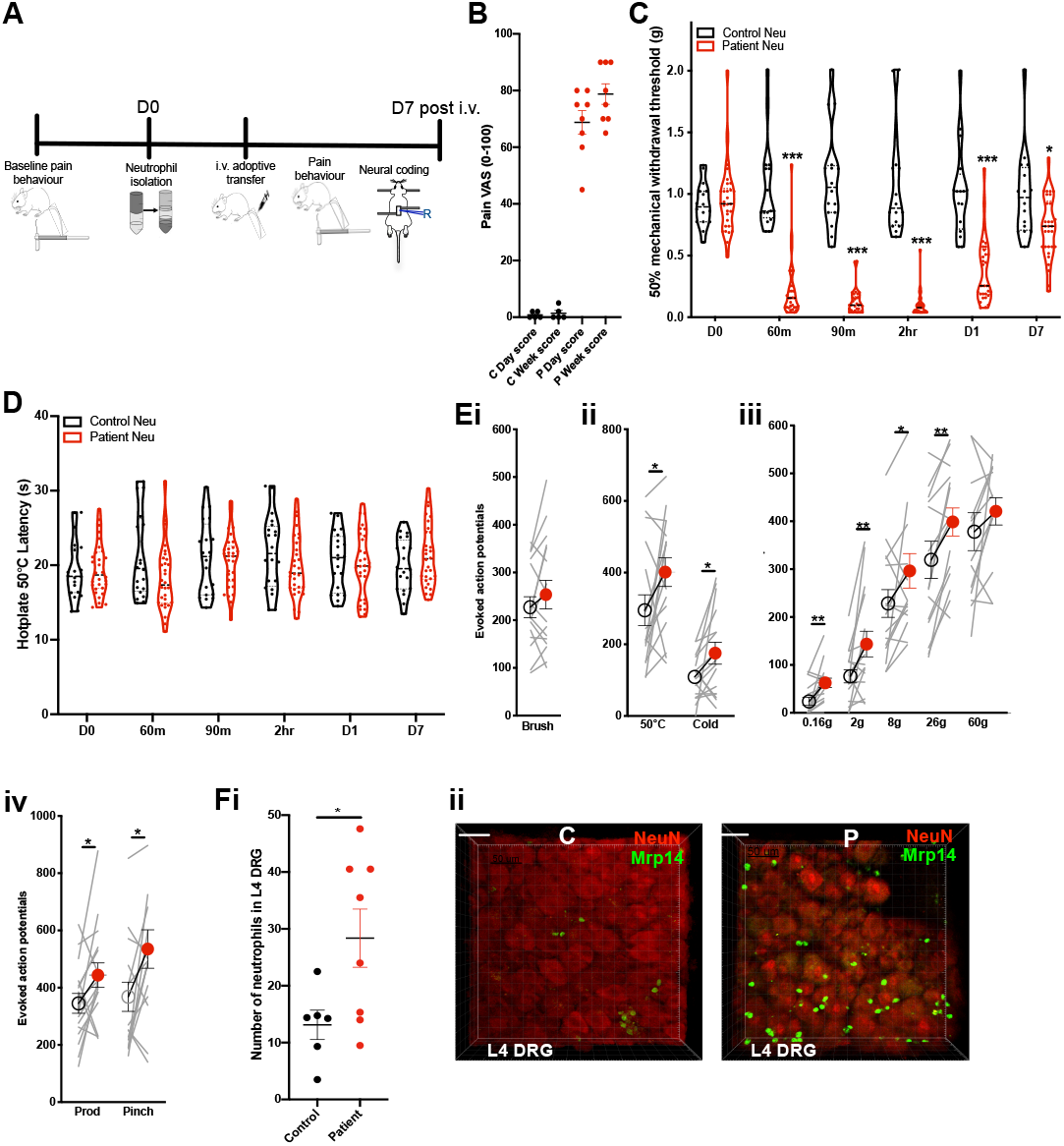
Adoptive transfer of neutrophils from primed mice confers widespread pain in naïve mice. **(A)** Neutrophils were isolated from patients with fibromyalgia (n=9) and healthy control subjects (n=5) and administered into naïve male and female mice (1×10^6^ cells i.v. in PBS suspension). Cells from each subject were injected into 4 mice. **(B)** Pain VAS scores of healthy control subjects and subjects with fibromyalgia. **(C)** Mechanical hypersensitivity to von Frey stimulation of mice following i.v. adoptive transfer of neutrophils (*P<0.05, ***P<0.001 2 way RM ANOVA Control vs Patient cells). **(D)** Latency to nociceptive behaviour in the hotplate assay (2 way RM ANOVA Primed vs Control). **(E)** Evoked action potential firing of neurons deep dorsal horn of naïve mice (n=16 cells) before (open circles) and 1hr after (red circles) administration of neutrophils from patients. Total number of action potentials fired over 10s stimulation with (i) innocuous brush, (ii) noxious thermal stimuli, (iii) punctate von Frey stimuli and (iv) noxious mechanical stimuli (*P<0.05, **P<0.01 Paired T-test). **(Fi)** *Ex vivo* imaging of neutrophil cell counts in L4 DRG following adoptive transfer of neutrophils from Patient (n=4 mice) or Control blood (n=3 mice). White line illustrates 50µm. (*P<0.05 Unpaired T-test). (ii) Representative of mean images for (Fi).

## Discussion

In the last decade of the 20^th^ century the first theories of a neurological basis for fibromyalgia were put forward and fibromyalgia syndrome has since largely been considered a disease of the central nervous system (24). We used a mouse model of chronic widespread pain and a back-translational adoptive transfer model of neutrophils from patients with fibromyalgia syndrome to demonstrate that neutrophils are peripheral pathological drivers of chronic widespread pain. Our data reveals 3 main findings: (1) a hyperalgesic priming paradigm can be used to induce chronic widespread pain in mice, which reproduces the bilateral sensory dysfunction and mechanical hypersensitivity observed in patients with fibromyalgia syndrome; (2) circulating neutrophils in mice and in patients with chronic pain are pro-nociceptive; (3) systemic neutrophil sequestration can prevent the development of chronic widespread pain without affecting homeostatic resolution of inflammation following injury.

### A murine hyperalgesic priming model induces chronic widespread pain

We used a hyperalgesic priming paradigm to induce persistent and widespread pain in mice in line with other studies that have used priming of nociceptive pathways to precipitate latent, bilateral hypersensitivity and to model the transition from acute to chronic pain (21-23, 25). Consecutive intramuscular injections of the acute inflammatory stimulus carrageenan, induced prolonged ipsilateral, mechanical hypersensitivity in mice and a spatiotemporal spread of hypersensitivity to the contralateral limb lasting more than 4 weeks (23) (Figure 1B). Intramuscular carrageenan induced mechanical hypersensitivity in both primed and control mice that resolved by 4d. Sensitisation to von Frey stimulation of the hind paw indicates the development of secondary hypersensitivity, beyond the primary site of injury in the gastrocnemius muscle. Further electrophysiological characterisation of neural coding confirmed sensitisation of ipsilateral dorsal horn neurones with receptive fields in the hind paw as a mechanism underlying this secondary hyperalgesia (26, 27).

Moreover, we observed behavioural and neuronal hypersensitivity specifically to mechanical, but not thermal stimuli. Previous studies have also reported bilateral mechanical hypersensitivity with a single intramuscular injection of carrageenan (28) or acidic saline (29), as well as with consecutive injections of intramuscular carrageenan (30). Intramuscular injections of carrageenan also produced significant weightbearing asymmetry in control and primed mice. Primed mice continued to display substantial weightbearing asymmetry following the second carrageenan administration until ca. 2 weeks, corresponding to the development of contralateral mechanical hypersensitivity (Figure 1B, 1C). Contralateral dorsal horn neurones were also sensitised to mechanical stimuli indicating either top-down bilateral modulation of spinal excitability by supraspinal centres (31) or a systemic aetiology for the development of bilateral hypersensitivity. We tested the hypothesis for the latter narrative, and observed that transfer of blood cells, but not serum, from primed mice confers transient mechanical hypersensitivity to recipient naïve mice (Figure 2).

### Adoptive transfer of neutrophils confers neuronal sensitisation and pain

Neutrophils are primary mediators of rapid innate host defense prior to the complex humoral and lymphocyte cellular processes of acquired immunity (32). The development of severe immune deficiency following iatrogenic neutropenia from cancer-related chemotherapy highlights the importance of neutrophils in the immune response to infection and injury (33). Some studies suggest an additional role for neutrophils in anti-nociceptive activity based on their atypical expression of opioid peptides (34, 35). In contrast, we observed that transfer of neutrophils from mice or patients with chronic widespread pain can induce transient hypersensitivity in otherwise healthy mice (Figure 2 and Figure 5). Moreover, antibody depletion of neutrophils can prevent the development of persistent mechanical hypersensitivity in mice with chronic widespread pain induced by injections of intramuscular carrageenan (Figure 3). The role of neutrophils in regulating acute inflammatory pain and resolution of inflammation is well established, but neutrophil-mediated modulation of nociceptive processing beyond the inflammatory resolution phase has not been explored (36, 37). Several other findings in animal studies point towards a pro-nociceptive role of neutrophils, ever since the first report of pro-algesic effects of articular neutrophils in joints of dogs, promoted by administration of lipopolysaccharides (38). Subsequent studies by Levine *et al*. showed that intraplantar administration of LTB_4_ and the complement factor 5a in the rat paw produces mechanical hypersensitivity that is dependent on neutrophil migration (39, 40). In addition, thermal hypersensitivity induced by nerve growth factor is inhibited in neutrophil-depleted rats (41), and migrating neutrophils during carrageenan-induced inflammation contribute to mechanical hypersensitivity partly by release of PGE_2_ (42).

We used a back-translational paradigm to demonstrate that circulating neutrophils in patients with fibromyalgia syndrome can sensitise sensory neurons and elicit behavioural hypersensitivity in mice. Recent work indicates an immunological basis for fibromyalgia symptoms as passive IgG transfer from patients to mice can confer mechanical pain behaviour (43), although we were unable to reproduce these findings (Figure S4). This may be due to differences in pain severity (measured through pain VAS) of the patient cohorts blood samples were derived from, and reflect the potential heterogeneity of disease pathogenesis among patients with fibromyalgia. In support of our findings, previous studies in humans demonstrate that recruitment of polymorphonuclear leukocytes by the potent chemoattractant LTB_4_ decreases pain thresholds, causing robust hyperalgesia in healthy subjects (44). Moreover, intraarticular steroids not only reduce neutrophil infiltration of knee joints in patients with osteoarthritis and inflammatory arthritis, but also reduce pain scores (45). The close correlation between reduction in pain VAS and neutrophil migration supports a causal link between neutrophil migration and nociception. Responses in the contralateral knee indicate a systemic effect of neutrophil migration on perceived nociception (45).

We did not observe any changes in thermally-evoked pain behaviours in the 50°C hotplate following administration of neutrophils from mice or patients with chronic widespread pain (Figure 2E and Figure 5D), but did observe marked sensitisation of spinal neurons to 50°C stimulation of peripheral receptive fields in the hindpaw. The discrepancy between integrated behavioural responses and the excitability of individual spinal cord neurons may relate to the recruitment of segmental inhibition of spinally-mediated reflexes with suprathreshold stimulation following contact of all 4 hindpaws on the hotplate (46-48). It is possible that pain behaviours evoked by unilateral thermal stimulation of the hindpaw could be sensitised by neutrophil administration. Previous findings have observed that inhibition of neutrophil migration with selectin inhibitor fucoidin can reduce thermally-evoked pain behaviours on the 51°C hotplate following ovalbumin administration (49). Moreover, antibody depletion of circulating neutrophils significantly reduces endoneurium infiltration and attenuates the induction of thermal hyperalgesia induced by partial peripheral nerve injury (50).

### De novo infiltration of neutrophils into sensory ganglia leads to chronic widespread pain

Immune cell infiltration into nervous tissue beyond primary sites of injury has previously been observed in chronic pain models and indicated in pro-nociceptive processes (17-19, 51). T-cell and neutrophil infiltration in dorsal root ganglia in mice with diabetic peripheral neuropathy accompanies tonic pain (19). In the chronic constriction injury model of neuropathic pain, de novo neutrophil infiltration of ipsilateral dorsal root ganglia is observed with direct contact between polymorphonuclear granulocytes and neurons, despite no change in local expression of cytokine-induced neutrophil chemoattractant-1 (18). Moreover, nerve injury in mice leads to T-cell and neutrophil infiltration of dorsal root ganglia and development of mechanical hypersensitivity that can be attenuated with inhibition of leukocyte-derived elastase (15, 52).

We observed *de novo* infiltration of neutrophils into lumbar DRG in primed mice that could be prevented with antibody depletion of circulating neutrophils (Figure 3). We also observed increased infiltration endogenous mouse and exogenous human neutrophils into mouse sensory ganglia following adoptive transfer of neutrophils from a patient with fibromyalgia (Figure 5A-B, 5F, S3). The chemo-attractant mechanisms underlying neutrophil infiltration into the sensory ganglia is likely related to our observation of an increased prevalence of subpopulations of circulating neutrophils with high expression of surface markers CD62L (L-selectin) and CXCR2 (receptor for IL-8 and other chemokines) in Primed mice, whereas following infiltration into the DRG of Primed mice there is a greater prevalence of neutrophil subpopulations with lower CD62L and CXCR2 expression (Figure 4C-H), typical of an activated and migrated phenotype (53, 54). These observations are supported by previous work demonstrating increased L-selectin expression of neutrophils in patients with fibromyalgia syndrome, suggesting a possible phenotypic population of neutrophils plays a role in chronic widespread pain (55). Other work in experimental autoimmune encephalomyelitis suggests that DRG infiltration of neutrophils can also be dependent on TLR-CXCL1 signalling (56). In line with these findings, adoptive transfer of myelin oligodendrocyte glycoprotein-stimulated neutrophils induces mechanical allodynia in recipient mice (57). The likely mechanism underlying neutrophil-mediated sensitisation of DRG neurons in our study is also unclear; several studies demonstrated that neutrophils release algesic mediators that activate and sensitise nociceptive afferents directly, or indirectly by eliciting the release of algesic mediators from other resident cells, including the leukotriene 8R,15S-diHETE (58), superoxide radicals (59) and cyclooxygenase-2 axcrtivation that triggers release of prostaglandins (59, 60). Nerve growth factor induced hyperalgesia is also dependent on circulating neutrophils (41, 61). Due to the short half-life of neutrophils (3-7 days) it is likely that that these immune cells harbour a reactive capacity for phenotypic changes in chronic pain states. Other studies have shown an extension of neutrophil lifespans in pathological states (62, 63). These studies support our finding that neutrophils – when derived either from mice with persistent widespread pain last more than 2 weeks or from patients with suffering from fibromyalgia pain over years – can undergo phenotypic changes enable and sustain sensitisation of peripheral sensory neurons.

## Conclusion

We used a model of hyperalgesic priming to induce chronic widespread pain in mice that resulted in robust and persistent ipsilateral and contralateral behavioural hypersensitivity and sensitisation of spinal cord neurons. Adoptive transfer of neutrophils from primed mice and from patients with fibromyalgia syndrome confers mechanical pain to recipient naïve mice, sensitises evoked action potential firing of spinal cord neurons and causes neutrophil infiltration into dorsal root ganglia. These data demonstrate that neutrophils are fundamental for the development of chronic widespread pain through infiltration of peripheral sensory ganglia. Further studies characterising the neutrophil phenotype in fibromyalgia syndrome may shed light on mediators of the cross-talk between these polymorphonuclear granulocytes and sensory neurons. Our findings suggest that targeting neutrophils may be useful therapeutic targets for pain control in fibromyalgia.

## Acknowledgements

This work was supported by Versus Arthritis UK (21734), Barts Charity (MGU0532), Wellcome Trust and Rosetrees Trust. The authors would like to thank Professor Mauro Perretti and Dr Suchita Nadkarni for their invaluable advice and discussions in the preparation of this manuscript.

## Methods

### Animals

All animal assays performed in this study were approved by the United Kingdom Home Office Animals (Scientific Procedures) Act 1986. C57Bl/6J male and female mice aged 8–10 weeks were kept on a 12-hour light/dark cycle and maintained under standard conditions (21±1°C, food and water *ad libitum*).

### Mouse behaviour

All behavioural experiments were performed by an experimenter blind to randomised allocation of mice to experimental groups. Mechanical hypersensitivity was assessed using the up-down von Frey method as described previously (64). Thermal nociceptive thresholds were determined by measuring latencies to nociceptive behaviour on a 50°C hot plate (Ugo Basile). Hindlimb weightbearing was measured using the incapacitance tester (Bioseb).

### Murine model of chronic widespread pain

A model of hyperalgesic priming was used to induce chronic widespread pain in mice with consecutive injections of carrageenan into the right gastrocnemius muscle 4 days apart (i.m. 3% 30μl, Sigma Aldrich). Control mice only received one injection on D0 (30).

### In vivo electrophysiology

Electrophysiological recordings were performed by an experimenter blind to experimental groups. Briefly, mice were anaesthetized with isofluorane (4%; 0.5L/min N_2_O and 1.5 L/min O_2_) and secured in a stereotaxic frame. Anaesthesia was reduced and maintained at 1.5% isoflurane for the remaining duration of the experiment. A laminectomy was performed to expose L3–L5 segments of the spinal cord and extracellular recordings were made from wide dynamic range neurones in the deep dorsal horn (lamina III–V, 200–600□μm) using parylene-coated tungsten electrodes (A-M Systems, USA. Mechanical and thermal stimuli were applied to the peripheral receptive field of spinal neurons on the hindpaw glabrous skin and the evoked activity of neurons was visualised on an oscilloscope and discriminated on a spike amplitude and waveform basis using a CED 1401 interface coupled to Spike2 software (Cambridge Electronic Design, UK). Natural stimuli (dynamic brush, vF hairs 0.16g - 60g, noxious prod 100gcm^−2^ mechanical stimulation; thermal water jet 35 - 50°C) were applied in ascending order of intensity to receptive fields for 10s and the total number of evoked spikes recorded.

### Neutrophil depletion

Neutrophil depletion was performed by i.v. injection of 50µg anti-Ly6G antibody (clone 1A8 Biolegend #127601) or PBS on Day 3 (24 hours prior to the second injection in the primed group).

### Patient samples

Blood samples were obtained from healthy, pain free control subjects or from patients diagnosed with fibromyalgia according to using ACR 2016 criteria (65) at Mile End Hospital, Barts NHS Trust. All subjects were asked to rate their overall pain score on a visual analogue scale from 0-100 (where 0 = no pain, 100 = maximum pain possible) for the day and over the past week. Healthy control subjects had pain scores of 5 or less. All subjects provided written informed consent, and samples were collected in compliance with HRA approval (REC 17/WS/0172, IRAS 60271). Blood samples were collected immediately after giving consent and record of pain VAS scores. Neutrophils were isolated by density gradient centrifugation at room temperature and red blood cells were removed using ACK lysing buffer. Naïve C57B/6 mice were injected with 1×10^6^ neutrophils (i.v. 100μl in PBS) from either patients or controls.

### Adoptive transfer whole blood components

Whole blood was collected from Control or Primed mice through terminal cardiac bleed. For blood cell isolation, ACK lysing buffer was used to remove red blood cells from heparin-treated blood, the sample of centrifuged at 300 x g for 5 minutes at RT and the supernatant was discarded. The pellet was resuspended in 5mL DMEM solution (Gibco) and centrifuged at 300 x g for 5 minutes at 4°C. The supernatant was discarded and was used to isolate either serum or blood cells for adoptive transfer into naïve recipient mice. injected into naïve mice (i.v. 100μl). Neutrophil (Ly6C+Ly6G+), T-cell (CD3+), B-cell (CD19+) and monocyte (Ly6C+Ly6G-) populations of cells were isolated from either primed or control mouse blood using FACS cell sorting and injected into naïve recipient animals (i.v. 1×10^6^ cells in 100μl PBS). IgG from whole blood was purified using protein A Sepharose beads (Merck). Serum was diluted 1:2 in PBS, passed through a protein A column, and the bound IgG was eluted using Ab Buffer Kit (Merck), and then the eluate was dialyzed overnight at 4°C in PBS using a 10 kDa dialysis membrane (Thermo Fisher Scientific). The concentration of IgG present after dialysis was determined using a BCA assay and adjusted by dilution with PBS. The IgG solution was stored at 4°C until i.p. administration (8mg) into naïve mice.

### Flow Cytometry

Flow-cytometry analysis was performed on mouse blood cells. Blood samples from mice were collected in 1.5ml Eppendorf tubes with 50mM EDTA and kept on ice. Red blood cells were removed by incubating the blood samples with 2mL Ammonium-Chloride-Potassium Lysing Buffer (Thermo Fisher Scientific #A1049201) for 3 to 5minutes at RT. Samples were centrifuged at 300 × g for 5 minutes at room temperature and the supernatant was discarded. This step was repeated once. The cell pellet was then gently resuspended, followed by the addition of 1mL ice-cold DMEM media and centrifugation at 300 x g for 5 minutes at 4°C. The supernatant was discarded, and the pellet was resuspended in 500μL Hanks’ Balanced Salt solution. Cells were stained using Zombie Aqua Live/Dead kit (Biolegend #423101) for 15 min, washed and incubated for 10 min with human Fc TruStain FcX (Biolegend #101320). Cells were stained for surface antigens. Antibodies used are listed in Table 1. All samples were acquired using a BD Fortessa and analyzed with FlowJo software (FlowJo, LLC).

**Table 1.** FACS mouse antibodies.

| antibody | clone | Fluorochrome | company | catalogue number | batch | dilution |
| --- | --- | --- | --- | --- | --- | --- |
| CD45 | 30-F11 | BV785 | Biolegend | 103149 | B263597 | 1 in 800 |
| CD11b | M1/70 | APC | Biolegend | 101212 | B269209 | 1 in 200 |
| Ly6G | 1A8 | APC/cy7 | Biolegend | 127624 | B348489 | 1 in 1600 |
| Ly6C | HK1.4 | A488 | Biolegend | 128022 | B304228 | 1 in 1600 |
| CD62L | MEL 14 | BV605 | Biolegend | 104438 | B266851 | 1 in 800 |
| CXCR2 | SA045E1 | PerCP/Cy5.5 | Biolegend | 149606 | B273375 | 1 in 400 |

### tSNE Analysis

tSNE analyses were performed with FlowJoTM version 10 software(FlowJo LLC), with Ly6G Ly6C double positive neutrophils. All Ly6G/Ly6C double positive neutrophils were merged with the concatenate tool and barcoded to track individual cells and to distinguish the condition. Finally, tSNE analyses were performed with the CXCR2, CD62L, CD11b, Ly6G andLy6C markers. The following settings were used: learning configuration: auto (opt-SNE), KNN algorithm: Exact (vantage point tree) and gradient algorithm: Barnes-Hut.

### Immunohistochemistry

Mice were anesthetised by overdose with pentobarbital and perfused transcardially with phosphate buffered saline (PBS) followed by 4% paraformaldehyde in 1X PBS. DRGs were dissected and fixed in 4% PFA for 2h, followed by 30% sucrose with 0.02% azide in 1X PBS. DRGs were placed in OCT and cut on a cryostat at 10μm, followed by mounting on SuperFrost Plus glass slides for staining. Infiltration of neutrophils into the DRG neurones in Naïve, Control and Primed mice was evaluated using immunohistochemical staining of S100A9 co-stained with NeuN, a marker for neurones. The tissue sections were blocked with 10% normal goat serum (Vector Labs S-1000) in PBSTx (0.3% Triton-X in 1×PBS). Sections were co-stained with a monoclonal rat anti-S100A9 (Abcam; ab105472) and recombinant rabbit anti-NeuN antibody (Abcam; ab177487) and their respective secondary antibodies (Alexa Fluor 594 anti-rabbit IgG, ThermoFisher; A11037 and Alexa Fluor 488 goat anti-rat, Abcam; ab150157). Sections were mounted and coverslipped with Vectashield Hardset Antifade Mounting Medium with DAPI (Vectorlabs; H-1500) and imaged using an Olympus fluorescent microscope. The total numbers of S100A9+ cells neurons were quantified in three DRG sections per slide and pooled per mouse using ImageJ. Neutrophils within 250μm of the tissue periphery were counted by a blinded experimenter using ImageJ.

### Wholemount ex vivo multi-photon microscopy

Mice were culled with isoflurane overdose followed by immediate harvesting of lumbar DRG, which were then incubated for 4h in 4% PFA at 4°C followed by incubation with perm block solution overnight (20% serum, 0.5% Triton X-100 and 10 x PBS). The following day, DRG were incubated overnight at 4°C with primary antibodies against MRP14 (1:200; Abcam #ab105472), CD31-647 (1:200; Biolegend #102515) and NeuN (1:200; Abcam #ab177487). Anti-MRP14 and anti-NeuN were conjugated with A488 and A555 respectively, carried out using labelling kits (Thermo Fisher Scientific) according to the manufacturer’s instructions. Samples were prepared for imaging by PBS immersion and covered with a coverslip. DRGs were observed with a Leica SP8 DIVE multiphoton confocal microscope (Leica Microsystems) equipped with a 25× 1.0 NA WI IR objective lens and a pulsed infrared laser. Most experiments were performed at 795nm and 1045nm (MP laser) excitation with an intensity between 17.2% and 25%, respectively. Images were acquired with a 0.53-um z step size with approximate z depth of 130μm. Pre-defined settings for laser power and detector gain (speed 8000, pixel size 346.32nm^2^) were used for all experiments. Three-dimensional Images were then analysed off-line using IMARIS software (Bitplane, Switzerland) and the number of neutrophils per field of view was quantified by creating isosurfaces on the neutrophil channels.

### Statistical analysis

All behaviour, electrophysiology and imaging studies were conducted by experimenters blinded to treatment. Statistical analyses were performed using Prism9 and R software. Behaviour studies were analysed using 2 way ANOVA with repeated measures and Bonferroni posthoc tests to compare experimental groups. Unpaired t-Tests were used for electrophysiology data and FACS events comparing experimental groups. Immunohistochemistry analyses involved a minimum of 3 DRG sections from at least 3 animals per experimental group using a one-way ANOVA.

## Supplementary figures

S1. Motor function of Control (*n*=7) and Primed (*n*=7) mice assessed on he rotaros demonstrating no differences between groups in (A) rotations per minute, (B) latency to fall and (C) distance travelled (P>0.05 for all time points 2 way RM ANOVA Primed vs Control).

S2. (i-ii) tSNE analysis of Ly6G/Ly6C double positive cells derived from (i) blood and (ii) DRG (of both Control and Primed mice) illustrating heterogenous expression of CXCR2, CD62L, CD11b, Ly6G and Ly6C. (iii) tSNE analysis of Ly6C/Ly6G double positive neutrophils obtained from blood (*n*=8 mice) and DRG (*n*=6 mice) of Primed animals. Differential expression of CXCR2, CD62L, CD11b, Ly6G and Ly6C based on DRG specific clusters also illustrated.

S3. Representative image of L4 DRG 24hr following i.v. administration of Patient neutrophils, illustrating endogenous neutrophils (mouse MRP14; green) and exogenous neutrophils (CellTracer dye 555; blue) infiltrating peripheral mouse ganglia. Some neutrophils are indicated by white arrows.

S4. Passive transfer of 8mg IgG derived from a fibromyalgia patient *vs*. pain free control into naïve recipient mice (*n*=4 mice per group). No differences in sensory thresholds were observed over a 7-day period to (i) punctate mechanical stimulation or (ii) noxious heat stimulation (P>0.05 for all time points 2 way RM ANOVA).

