## Supplementary figures and images for "Neutrophils infiltrate sensory ganglia and mediate chronic widespread pain in fibromyalgia"

### S1

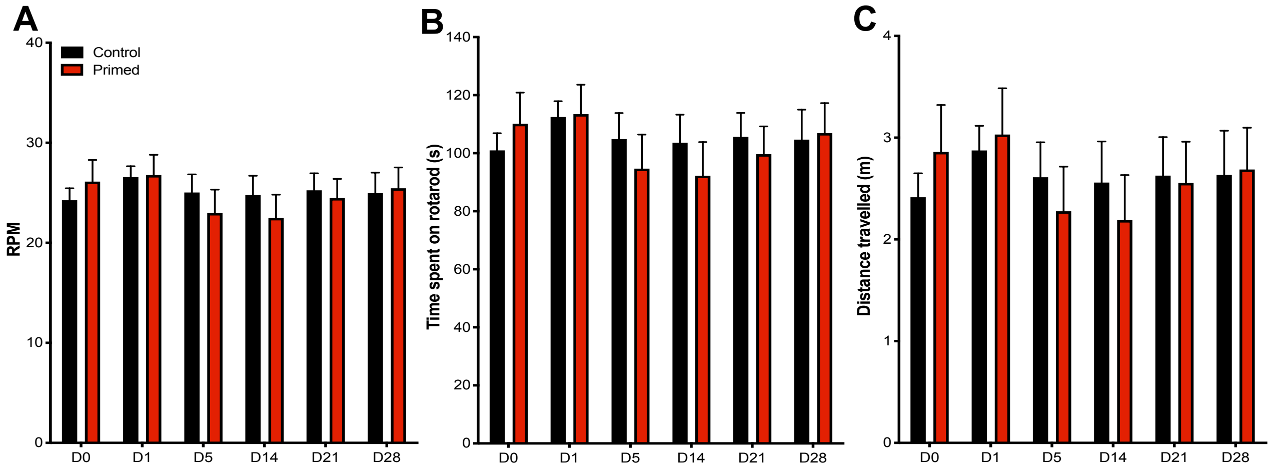

### S1

S2i

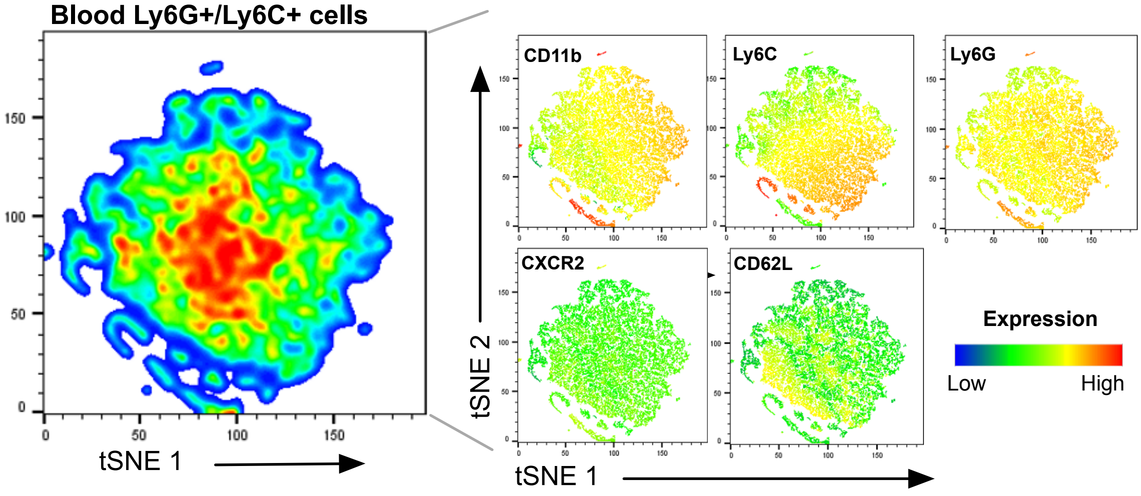


S2ii

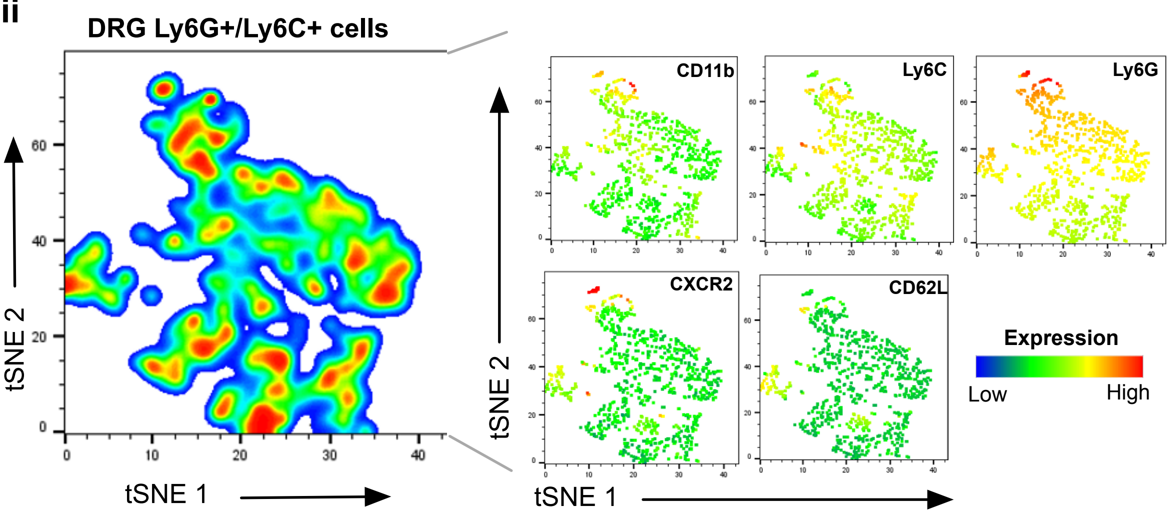


S2iii

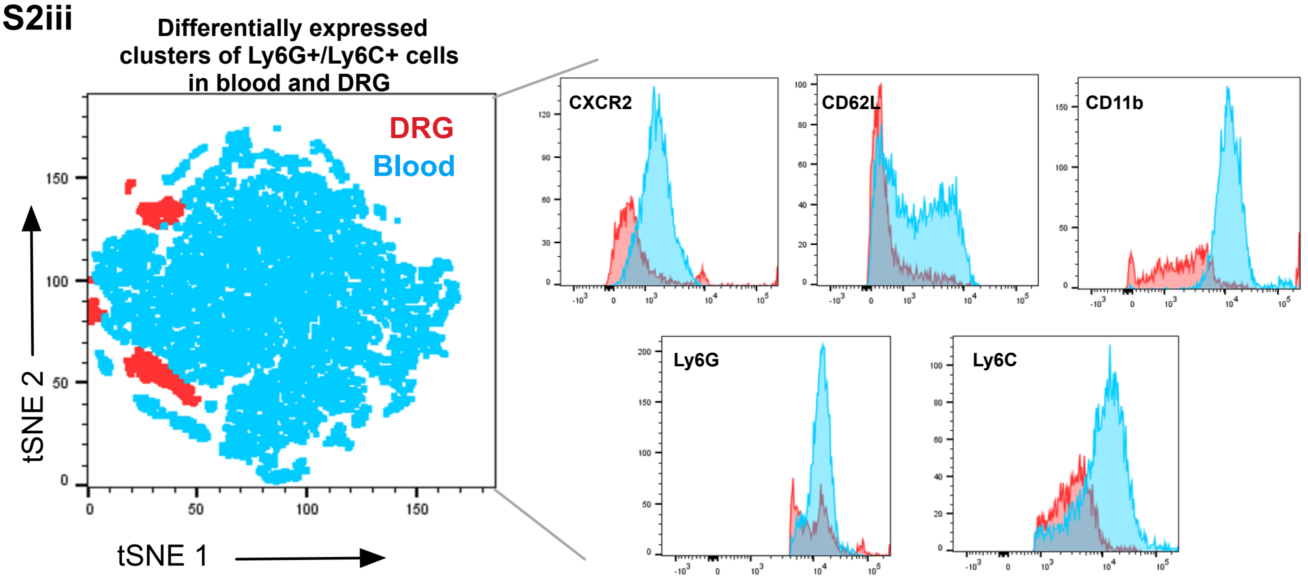

### S3

**S3**

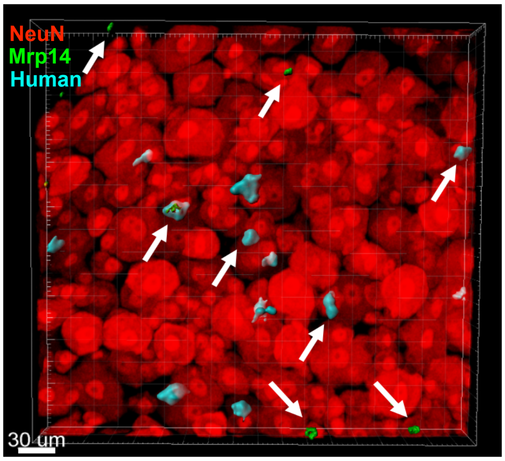

### S4

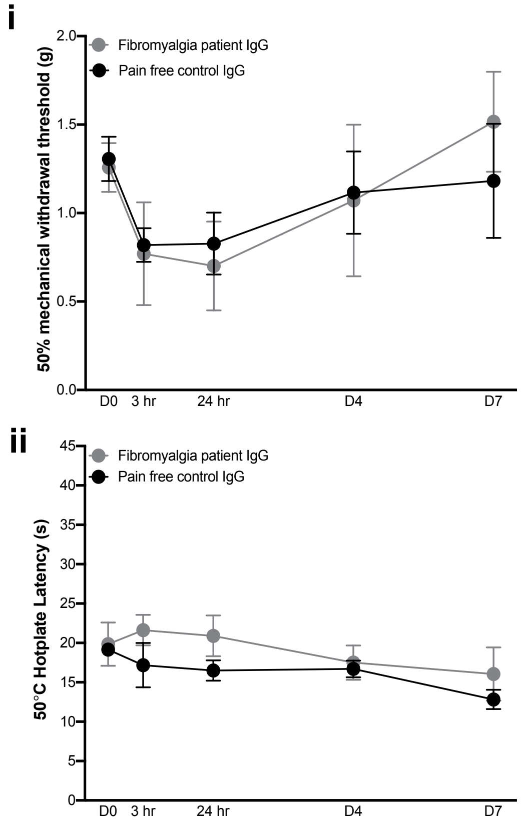
